# Macrophages of diabetes and tuberculosis co-morbid conditions show perturbed differentiation and metabolic signatures

**DOI:** 10.1101/2025.03.05.640816

**Authors:** Nidhi Yadav, Nikhil Bhalla, Ashish Gupta, Shweta Chaudhary, Ranjan Kumar Nanda

**Author notes:** **Corresponding author** Ranjan Kumar Nanda (PhD), Translational Health Group, International Centre for Genetic Engineering and Biotechnology, New Delhi-110067, India, **E. mail:**.

## Abstract

Diabetes negatively impacts the host immune system, leading to allergies, recurring infections, and lameness. Its impact on the distribution and molecular phenotypes of the bone marrow-derived macrophages (BMDMs) needs attention and, more importantly, in co-morbid conditions like diabetes and tuberculosis (DM-TB).

In this study, total bone marrow cells were harvested at 3 w.p.i., from the low aerosol dose (100-120 CFU) of *Mycobacterium tuberculosis* H37Rv infected control and NA-STZ induced diabetic C57BL/6 mice. The bone marrow cells were differentiated into BMDMs, and CD11c, CD11b and F4/80 marker expressions were monitored using flow cytometry. BMDMs stimulated with Mtb H37Rv lysate were subjected to global transcriptome profiling to identify the perturbed metabolic and molecular landscape.

The BMDMs of the DM-TB comorbid mice showed low CD11c, CD11b and higher F4/80 expression. The macrophages of the DM group showed a skewed pro-inflammatory response, characterized by perturbed complement response, NF-kB pathway and fatty acid metabolism, whereas macrophages of the DM-TB group showed deregulated ceramide and amino acid metabolism. Macrophages from the DM and DM-TB group stimulated with Mtb H37Rv lysate, showed a deregulated transcript signature that impacts humoral response.

The dampened immune response observed in the DM-TB group macrophages highlights the importance of identifying novel targets for developing tailored drugs for comorbid conditions.

## Introduction

Worldwide, an annual incidence of 537 million diabetes (DM) cases are reported in 2021 (Webber, 2013). This increase in DM epidemic is also impacting the worldwide tuberculosis (TB) case incident rates (Jeon & Murray, 2008). DM patients were reported to have three times higher susceptibility to TB development and higher death rates during treatment (Zheng et al., 2017). The DM-TB patients present with higher mycobacterial load and cavitary lesions, indicating a more severe form of disease (Chaudhary et al., 2025). Therefore, DM-TB patients need separate attention, and a better understanding of their dampened immune response at cellular and molecular levels is critical.

Pathologic stress in diabetic individuals leads to sustained pro-inflammatory cytokines in circulation. This prolonged increased cytokine level is termed meta-inflammation and leads to sensitization of the system and reduced cellular phagocytic ability (Pavlou et al., 2018). Metabolic imbalances such as hyperglycemia and dyslipidemia also induce epigenetic modifications that exacerbate macrophage activation and the production of pro-inflammatory mediators, further advancing disease progression (Witcoski Junior et al., 2024). The knowledge about metabolic reprogramming of macrophages observed in DM individuals is limited to tissue-resident and circulatory macrophages. The influence of diabetes on the monocytic precursors in the bone marrow microenvironment in detail is yet to be explored.

In this study, we aimed to identify key modulators that may explain the dampened immune response from bone marrow-derived macrophages (BMDMs) in DM conditions and which could be targeted for improved outcomes in DM-TB comorbid conditions. We focused on the cellular and molecular landscape of the BMDMs harvested from different conditions and how they get altered upon stimulation with *Mycobacterium tuberculosis* (Mtb) H37Rv lysate.

## Results

### BMDMs of eu- and hyper-glycemic mice show varied phenotypes

The chemical method of DM induction, adopted in this study, replicates the T2DM pathology (**Figure 1A**). The DM mice consistently showed hyperglycaemia (>300 dg/mL) and dyslipidaemia, and the results are presented in an earlier report. The DM and control mice were aerosol infected with low doses (100-120 cfu) of animal passaged Mtb H37Rv strain. At 3 w.p.i., total bone marrow cells were harvested from the Mtb infected and control groups, which showed similar viability (**Figure 1B**). However, the bone marrow of diabetic mice showed higher Mtb dissemination (50%/20%: DM-TB/TB) (**Figure 1C**). These bone marrow cells were differentiated using GM-CSF, and the DM-TB group yielded a higher number of macrophages than the TB group and similar between the HC and DM control groups (**Figure 1D**). This indicates that the bone marrow of DM-TB had higher proliferative and differentiation potential. Interestingly, the percentage of CD11c+ and CD11b+ population was significantly lower in the DM-TB group than in healthy controls, whereas the percentage of F4/80+ cells was similar (**Supplementary Figure 1**). The DM-TB BMDMs had significantly lower CD11c expression than the healthy control, whereas CD11b expression was significantly low in Mtb-infected TB and DM-TB (**Figure 1E**). Higher F4/80 expression was observed in these cells of the DM-TB group but was low in the TB groups compared to the healthy controls (**Figure 1E**). Our findings suggest altered immunophenotype and proliferative potential in DM-TB comorbid mice.

**Figure 1:**
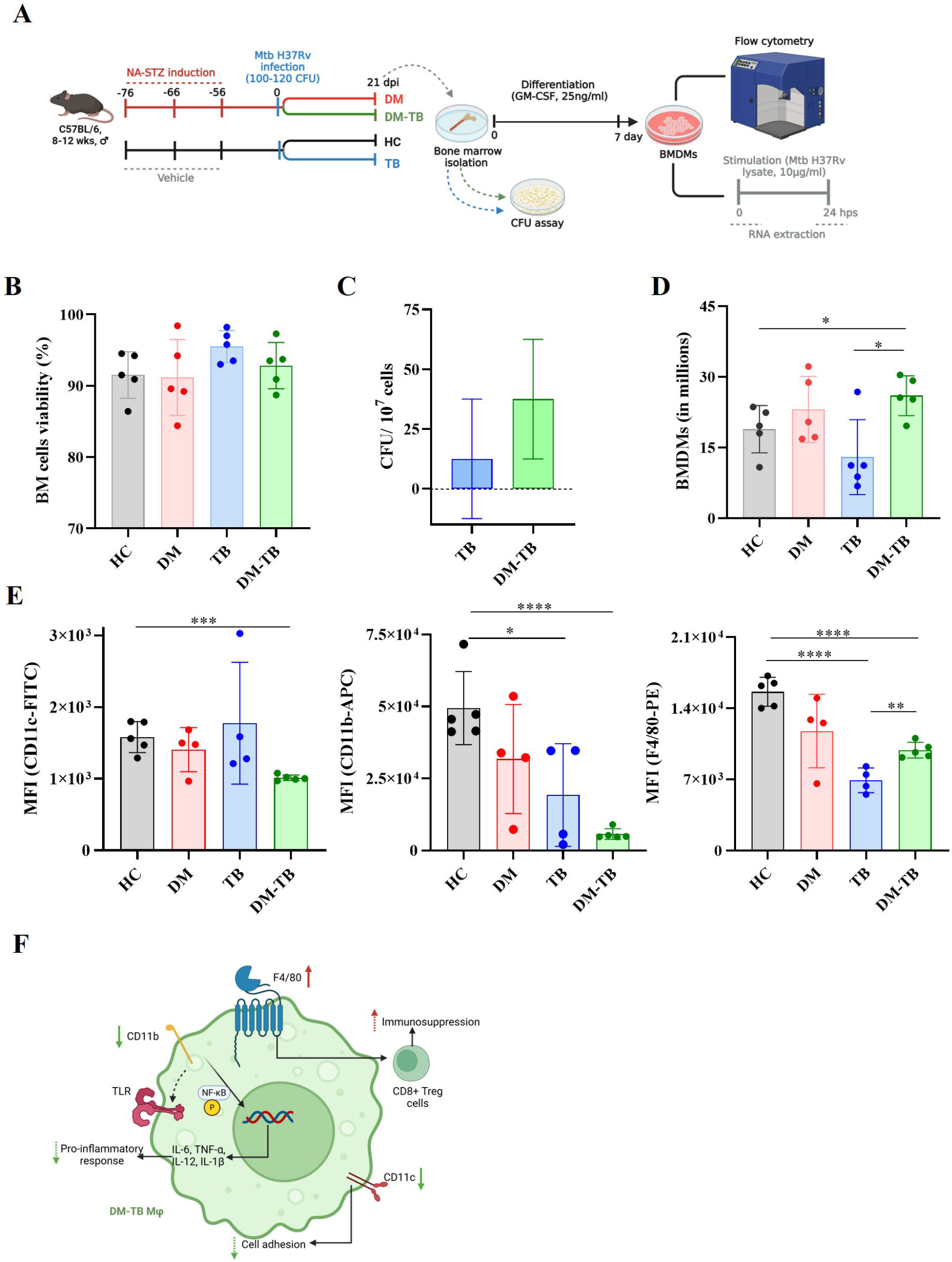
Bone marrow-derived macrophages from diabetes-tuberculosis comorbid mice show altered immunophenotype. **A**. Schematic presentation of the method used for diabetes induction and aerosol infection and bone marrow harvesting, differentiation and processing **B**. Viability of BM cells assessed by trypan blue assay **C**. Mycobacterial burden in the bone marrow of infected groups assessed by CFU assay **D**. Number of differentiated BMDMs obtained after differentiation **E**. Expression of CD11c, CD11b and F4/80 markers on live BMDMs **F**. Macrophages from DM-TB comorbid mice show lower CD11c, CD11b expression and higher F4/80 expression with further alteration in DM-TB comorbid condition which might decrease the pro-inflammatory response against pathogen. ns: not significant; *: p-value > 0.05; **: p-value > 0.005; ***: p-value > 0.001; BM: Bone marrow; BMDM: Bone marrow-derived macrophage; MFI: Mean fluorescence intensity; H: Healthy; DM: Diabetic; TB: Tuberculosis; DM-TB: Diabetic tuberculosis comorbid group; CFU: Colony forming unit; GM-CSF: Granulocyte monocyte-colony stimulating factor.

### Transcriptome profile of BMDMs harvested from the hyperglycemic mice showed a dampened immune response against Mtb

The transcriptome profile of the BMDMs harvested from the DM-TB group and controls were analysed to identify group-specific differences. Harvested total RNA, with an integrity >7.0 was subjected to Illumina sequencing to monitor hyperglycemia-associated changes in the immune response and macrophage metabolism (**Figure 2A and 3A**). The GC percentage of the RNA-seq data was 51.4±0.2%; the insert size in the alignment file was 310.5±6 bp; the alignment rate was 13.6±7.1%; 9.3±4.7 out of 67.8±1.8 million reads aligned to the reference genome. A total of 44,147 query mRNAs were assembled into 24,996 loci using StringTie, with an average of transcripts per locus. The reference annotation consisted of 43,518 mRNAs in 24,996 loci, and the number of super-loci containing reference transcripts was 24,996. A set of 39,508 matching intron chains, 43,256 matching transcripts, and 24,865 matching loci were identified. All 44,147 consensus transcripts were written to the merged annotated GTF file, with none discarded as redundant.

**Figure 2:**
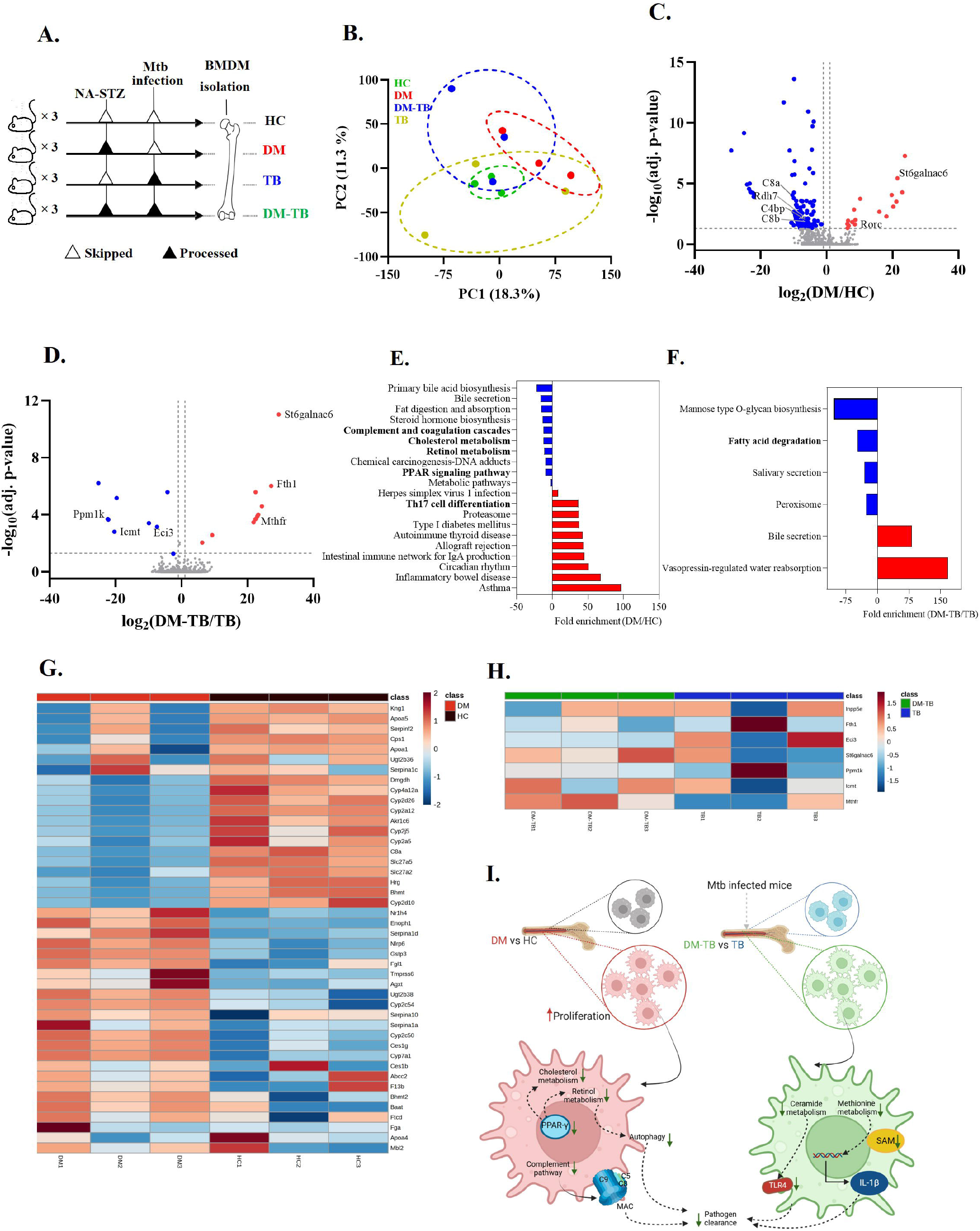
Diabetes induces transcription changes in bone marrow cells, which alters the macrophage metabolism, leading to a dampened immune response against Mtb. **A**. Schematic presentation of the experiment. **B**. Principal component analysis of BMDM gene expression data from various groups **C** and **D:** Pairwise transcriptome comparisons in the form of volcano plots are shown. **E** and **F:** Pathway enrichment analysis. **G** and **H**. Heatmap showing genes associated with immune response and altered metabolism between DM vs HC and DM-TB vs TB respectively. **I. Dampened antibacterial response in macrophages from hyperglycemic mice:** BMDMs from diabetic mice showed increased proliferation, decreased retinol metabolism and complement pathway. The Complement pathway and retinol metabolism regulate the inflammatory response. Downregulation of these pathways might lead to dampened Membrane attack complex (MAC) formation, autophagy and hence decreased pathogen clearance. However, the DM-TB comorbid mice showed decreased ceramide and amino acid metabolism. Ceramide metabolism regulates TLR signaling leading to downregulated pathogen clearance. Downregulated methionine metabolism contributes to S-adenosyl methionine (SAM) production which helps in the transcription of cytokines in response to pathogens. Student’s T-test was performed to determine the genes with significant group differences. Normalized counts to Z-scores of the genes that showed P<0.05 were used for unsupervised clustering and heatmap. BMDM: Bone marrow-derived macrophages, Mtb: *Mycobacterium tuberculosis*, HC: Healthy controls, DM: Diabetic, TB: Tuberculosis, DM-TB: Diabetic-tuberculosis comorbid group

Principal component analysis (PCA) of the RNAseq data of the BMDMs harvested from the HC, DM, TB and DM-TB groups showed partial overlapping clusters (**Figure 2B**). A set of 137 deregulated genes (17/120: up/downregulated) were identified in the BMDMs of the hyper-glycemic group compared to the eu-glycemic controls. BMDMs of the hyper-glycemic condition showed a downregulated complement pathway with a 10-fold decrease in the C8a complement gene expression with other genes (C4bp, C5, C8b and C9) involved in forming membrane attack complex to fight bacterial invasion (**Figure 2C**). The metabolic state of the macrophages drives their inflammatory response. In the BMDMs of the diabetic mice, an altered retinol, steroid and cholesterol metabolism was observed (**Figure 2E**). Macrophages of the diabetic mice showed a downregulated retinol metabolism with more than fivefold downregulation of retinol and retinal dehydrogenase (Rdh7 and Aldh8a1, respectively). PPAR (peroxisome proliferator activator receptor) pathway was downregulated in the BMDMs of diabetic mice (**Figure 2E**).

To understand the role of hyperglycemia-associated changes against Mtb infection, the transcriptome profiles of the BMDMs from DM-TB and TB groups were compared. A set of 275 deregulated genes (140/135: up/downregulated) were identified in the BMDMs of DM-TB mice compared to TB controls (**Figure 2D**). Amino acid metabolism, especially cysteine metabolism in the BMDMs, was downregulated in DM-TB comorbid mice (**Figure 2F**). Mthfr (methylenetetrahydrofolate reductase), which is involved in converting 5,10-methylenetetrahydrofolate to 5-methyltetrahydrofolate, is important for the remethylation of homocysteine to methionine, showed >20-fold upregulation in the BMDMs of the DM-TB group. Interestingly, Icmt1 (isoprenylcysteine carboxyl methyltransferase), which adds a methyl group to the C-terminal cysteine residue of proteins, showed >20-fold downregulation (**Figure 2G**). BMDMs of the DM-TB group showed a downregulated fatty acid metabolism, primarily for the glycosylceramide metabolism and Protein phosphatase (Ppm1k), Icmt and Mthfr showed >20-fold downregulation, which may directly impair the one-carbon metabolism. These findings suggest that the BM precursors from different groups differentiated with similar media composition show group-specific variations, including compromised lipid metabolism macrophages in the DM-TB group, which might affect pathogen clearance.

### Mtb whole cell lysate stimulation perturbs the BMDMs transcriptome in a group-specific pattern

We carried out an additional transcriptomics experiment to identify the deregulated pathways in the BMDMs of the DM-TB group and controls after stimulation with Mtb H37Rv whole cell lysates (Figure 3A). In this experiment, the Mtb-infected group was excluded due to low-quality RNA yield. The PCA plot showed overlapping clusters in the transcriptome profiles of the BMDMs harvested from the DM, DM-TB and HC groups (**Figure 3B**). A set of 45 (22/23: up/downregulated) deregulated genes were identified in BMDMs of DM mice compared to the healthy group post-stimulation (**Figure 3C**). The deregulated genes in the macrophages of diabetic mice were related to the TLR or humoral response and amino acid metabolism. Reg3g, Rnf170, and Icmt genes showed >20-fold downregulation in the Mtb lysate stimulated macrophages of diabetic mice compared to the healthy mice (**Figure 3E**). The macrophages of the DM-TB group, when stimulated with Mtb lysate, responded differently, and a set of 176 (89/76: up/downregulated) deregulated genes were identified (**Figure 3D**). Interestingly, the antimicrobial activity via humoral response showed downregulation in the Mtb lysate-stimulated macrophages of the DM-TB group, whereas Gata1 showed five-fold upregulation (**Figure 3F, G**). These findings suggest that BMDMs from different groups show distinct responses to Mtb stimulation.

**Figure 3:**
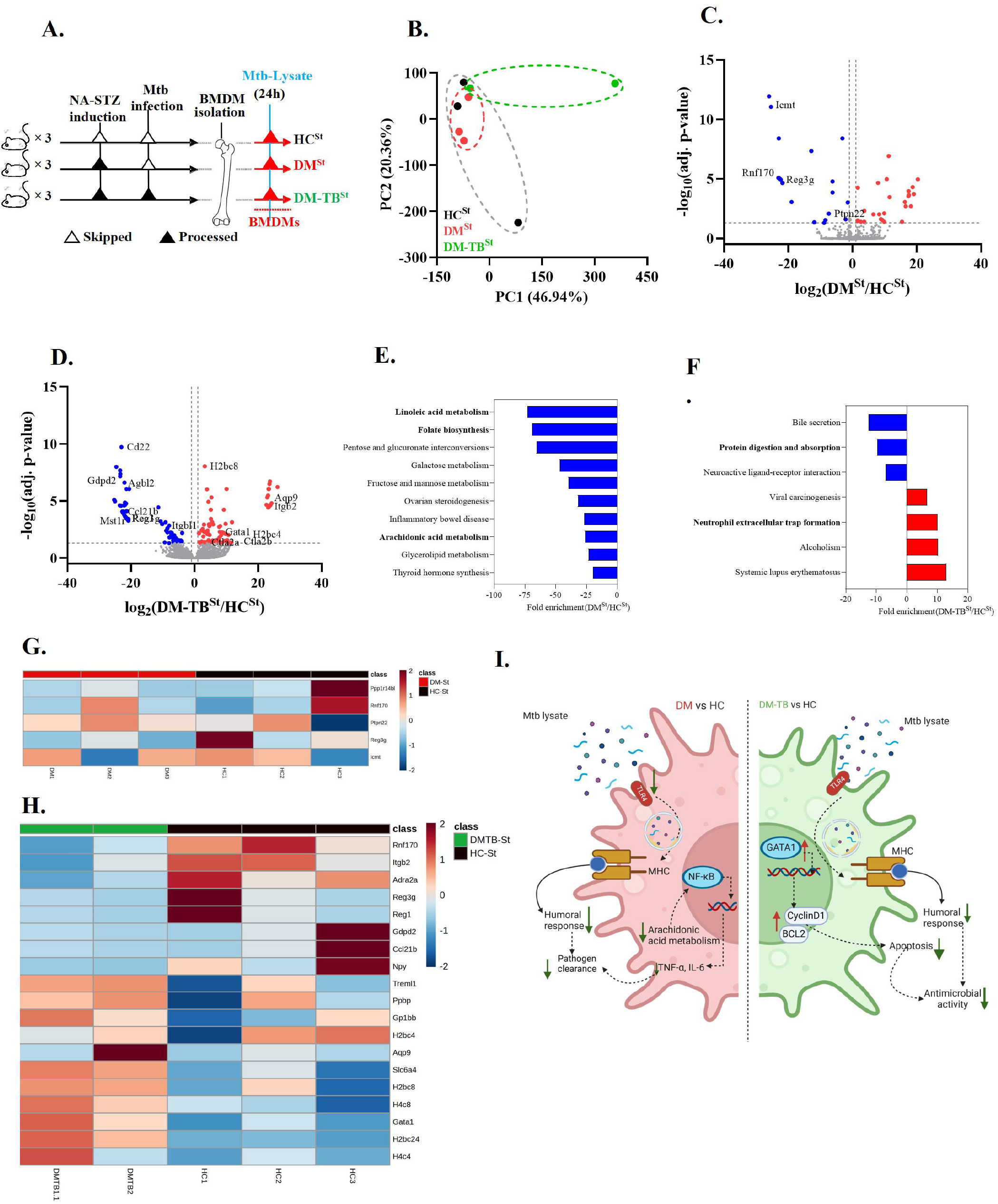
Diabetes-induced transcriptomic changes lead to altered macrophage metabolism leading to poor immune response on stimulation with Mtb lysate. **A**. Schematic presentation of the experiment. **B**. Principal component analysis of BMDM transcriptome **C** and **D**. Pairwise transcriptome comparisons of BMDMs from different sources stimulated with Mtb lysate in the form of volcano plots are shown. **E and F:** Pathway enrichment analysis plots **G** and **H**. Heatmap of deregulated genes involved in mentioned pathways in hyperglycemic mice BMDMs stimulated with Mtb lysates as compared to the euglycemic controls. **I. Macrophages from hyperglycemic mice respond differently to Mtb lysate:** BMDMs from DM and DM-TB mice respond differently to Mtb stimulation as compared to Healthy controls. Macrophages from DM and DM-TB group showed downregulated TLR4 signaling and humoral response. Arachidonic acid metabolism also got downregulated which will lead to dampened NF-kB signaling and inflammatory cytokine production. Macrophages from DM-TB group showed upregulated GATA6 which increases transcription of CyclinD1 and Bcl2 leading to decreased apoptosis and hence these cells could serve as niche for pathogens like Mtb. Student’s T-test was performed to determine the genes with significant group differences. Normalized counts to Z-scores of the genes that showed P<0.05 were used for unsupervised clustering and heatmap. BMDM: Bone marrow-derived macrophages, Mtb: *Mycobacterium tuberculosis*, DM: Diabetic, HC: Healthy controls, DM-TB: Diabetic tuberculosis comorbid group, ^St^: Stimulated with Mtb lysate

## Discussion

Macrophages belong to the innate arm of the immune system, representing the first line of defence against invading pathogens in vertebrates and invertebrates (Cooper & Alder, 2006). Depending on the stimulus or cytokine interacting with their surface or cytoplasmic receptors, macrophage populations show either pro- or anti-inflammatory phenotypes.

Hyperglycemia influences macrophages and sensitizes them to alter the cytokine production ability, leading to reduced phagocytic ability (Pavlou et al., 2018). In this study, eu- and hyperglycaemic C57BL/6 mice were infected with Mtb via aerosol mode, and the bone marrow cells were collected three weeks post-infection. Mtb is reported to disseminate to the bone marrow, and we observed that 50% of the DM-TB mice had Mtb in their bone marrow cells, whereas in euglycemic mice, Mtb reached to the bone marrow in only 20% of mice (Bobba et al., 2023). Interestingly, the overall monocyte (CD11b^+^CD11c^+^) numbers decreased in the bone marrow of the DM-TB group, and an increased F4/80 expression was observed. However, the proliferative capacity increased in terms of differentiated macrophage numbers. CD11b, an integrin protein, helps macrophages regulate TLR signalling in response to any intracellular pathogen and activation of NF-kB cascade and pro-inflammatory response (Yao et al., 2019). BMDMs of the DM-TB group had low CD11b expression, which might contribute to the altered phagocytic ability, as reported elsewhere (Ref). CD11c, an integrin alphaX transmembrane protein, a marker for innate immune cells, is involved in leukocyte adhesion and helps the macrophages in mounting immune response after reaching the site of infection. Low CD11c expression in the BMDMs of DM-TB mice might also impair the phagocytic ability. Low macrophage CD11c and CD11b expression alters leukocyte adhesion and TLR signalling. This may affect the NF-kB activation leading to low production of pro-inflammatory cytokines, impacting the response against intracellular pathogens (Yao et al., 2019, Arnold et al., 2016).

Along with decreased innate and monocytic marker expression in the BMDMs of the DM-TB mice, we observed higher expression of F4/80, a macrophage-specific adhesion GPCR. F4/80 is involved in IFN-γ production and immunoregulation. Increased F4/80 expression induces regulatory T cell production and increases immunosuppression; hence, the sensitized macrophages from the DM-TB group might show immunosuppressive behaviour upon encountering pathogens or pathogenic products (Lin et al., 2010) (**Figure 1F**).

Components of the complement pathway, including C4, C3 and factor D, are reported to be elevated in diabetes mellitus patients (Shim et al., 2020). Interestingly, the C8a complement gene, which forms a membrane attack complex to fight the bacterial invasion, was downregulated (∼10 fold) in the BMDMs from diabetic mice compared to the healthy controls. The complement pathway was downregulated in the BMDMs of the hyperglycaemic group. Other downregulated genes belonging to the complement pathway were C4bp, C5, C8b and C9 (**Figure 2C, G**). A dampened complement pathway is associated with poor opsonization of bacteria, affecting pathogen clearance (Jagatia, Heena and Tsolaki, 2021). This shows that even though diabetic individuals have high circulatory, endothelial and adipocyte-associated complement levels leading to chronic inflammatory state and macrovascular complications, the sensitized bone marrow cells have a decreased ability to produce the complement components, which would lead to lowered phagocytic ability.

Retinol metabolism is involved in macrophage polarisation (Acharya et al., 2022). BMDMs of DM mice showed more than fivefold downregulation of retinol and retinal dehydrogenase (Rdh7 and Aldh8a1, respectively). Rdh7 is involved in producing all-trans-retinol (atRA), which promotes autophagy and accelerates mycobacterial clearance (Chen et al., 2015). Aldh8a1 promotes Th2 response against pathogens, especially helminths and its downregulation in diabetic individuals might compromise their response to infections (Esposito et al., 2024). Retinoic acid (RA) is secreted by antigen-presenting cells, including macrophages, and helps in T-cell activation. RA signalling regulates inflammatory and regulatory macrophages and influences T cell functions. PPAR signalling also regulates the polarisation of macrophages and is needed for the effective clearance of apoptotic cells. It also helps macrophages switch from a pro-inflammatory (M1) phenotype towards an anti-inflammatory (M2) state (Abdalla et al., 2020). PPAR (peroxisome proliferator activator receptor) pathway was downregulated in the BMDMs of diabetic macrophages (**Figure 2C**). This sustained inflammation may not resolve over time due to the delayed switch to M2 polarization.

We monitored hyperglycemia-associated changes in metabolism, which may lead to distinct immune responses and pathophysiological outcomes in the DM-TB comorbid condition compared to the TB controls. We observed impaired amino acid and fatty acid metabolism in the DM-TB macrophages compared to the TB group. Mthfr (methylenetetrahydrofolate reductase) which converts 5,10-methylenetetrahydrofolate to 5-methyltetrahydrofolate, important for the remethylation of homocysteine to methionine, was upregulated whereas Icmt1 (isoprenylcysteine carboxyl methyltransferase) which adds methyl group to the C-terminal cysteine residue of proteins was downregulated in the DM-TB group (Raghubeer, Shanel and Matsha, 2021, Yang et al., 2020). This suggests that methyl groups from homocysteine are utilized in methionine formation rather than being used for protein modifications. Macrophages of the DM-TB group showed downregulated fatty acid metabolism, especially glycosylceramide metabolism. Ceramides contribute to proinflammatory response via TLR4 signalling (Mobarak et al., 2018). Protein phosphatase (Ppm1k), which maintains the repopulation capacity of bone marrow precursors, was downregulated in the DM-TB group. This implies deregulated branched-chain amino acid metabolism and impaired ability of bone marrow cells to proliferate in the tissues after infection (Li et al., 2023). Icmt and Mthfr were also downregulated, leading to impaired one-carbon metabolism and, hence, downregulated pro-inflammatory response against pathogens due to lowered S-adenosyl methionine production for methyl group transfer to histones. Deregulated cysteine-methionine metabolism is also reported to contribute to worsened liver pathology in DM-TB comorbid mice (Chaudhary et al., 2024).

Since hyperglycaemic individuals show impaired immune response, cytokine production and autophagy against pathogens, we also tried to monitor the transcriptomics differences in the BMDMs from diabetic mice after stimulating with Mtb H37Rv lysate. Our findings corroborate earlier transcriptome analysis of the macrophages isolated from DM condition, indicating an altered TLR response, humoral response and amino acid metabolism (Sousa et al., 2023). Downregulation of Reg3g, Rnf170, and Icmt genes was observed in the Mtb lysate-stimulated diabetic group compared to healthy mice macrophages. Reg3g shows bactericidal activity, and in chronic inflammatory conditions, higher expression of Icmt is associated with TLR-mediated macrophage activation via the Ras/MAPK pathway (Chakraborty et al., 2024). (Yang et al., 2020). Downregulation of these genes in the macrophages of the DM group indicates compromised activation and bactericidal activity.

It is reported that diabetes-tuberculosis comorbid individuals not only have increased mortality and worsened treatment outcomes but also show delayed adaptive immune priming (Prada-Medina et al., 2017). We compared the transcriptomics profile of the DM-TB comorbid group with the healthy controls post-Mtb stimulation. Antimicrobial activity via humoral response showed downregulation in the DM-TB group post-Mtb stimulation. However, the stimulated macrophages from the DM-TB group showed increased immune cell maturation. Gata1 is reported to block IL-6 induced macrophage differentiation and apoptosis through sustained expression of Bcl2 and cyclin D1 and was upregulated in the comorbid group post-stimulation.

We have uncovered the transcriptome differences in the BMDMs from hyperglycaemic and euglycemic mice after Mtb infection. Even though the bone marrow precursors were differentiated with similar media composition, still the macrophages of the DM and DM-TB groups showed distinct transcriptome profiles as compared to the controls. Whether these transcriptome-based differences would lead to a compromised pathogen clearance by altering cytokine levels is yet to be established.

In this study, we report that the altered immune maturation and humoral response in the diabetes group may need separate resolution and more importantly in the comorbid DM-TB conditions. The deregulated molecular pathways identified in this study explain the perturbed humoral response observed in DM-TB comorbid conditions and are useful to better understand pathology and design novel therapeutics.

## Methodology

### Ethical statement

The experiments performed in this study were approved by the Institute Biosafety and the Institute Animal Ethics Committee (vide reference ICGEB/IAEC/07032020/TH-13Ext) of the International Centre for Genetic Engineering and Biotechnology, New Delhi.

### Mice model for diabetes induction

The chemical method of diabetes induction was carried out using nicotinamide (NA) and streptozotocin (STZ), as explained in earlier reports (Chaudhary et al., 2025). Briefly, NA (60 mg/kg in 0.9% saline) was introduced 30 mins before the intraperitoneal introduction of STZ (150 mg/kg in 50 mM citric acid buffer) in 16 hours fasted 8-12 weeks old male C57BL/6 mice in three doses with a 10 days interval. The control group received only the vehicle. After the last NA-STZ dose, fasting (6 hr) blood glucose levels were monitored using a glucometer in all the mice groups. Detailed information about DM induction and the parameters are presented in an earlier report (Chaudhary et al., 2025). After two months of hyperglycemia in DM mice were subjected to a low aerosol dose of Mtb H37Rv infection with the control.

### Mtb H37Rv culture and whole cell lysate preparation

Mtb H37Rv strain was revived from glycerol stocks stored at -80 ºC by inoculating in 7H9 media (BD Difco Middlebrook) supplemented with OADC (10%, Oleic acid, albumin, dextrose and catalase), glycerol (0.2%), tween-80 (0.05%) till mid-log phase (OD ∼ 0.6). The PDIM levels of the Mtb H37Rv used for mice infection experiments were monitored. Briefly, an aliquot of H37Rv bacterial suspension used for infection was cultured in a conical flask till the log phase and centrifuged at 3,500 rcf at room temperature for 10 minutes. The pellet was incubated with a mixture of methanol and chloroform (15 ml, 2:1) for 48 hours to collect the supernatant by centrifuging at 3500 rcf for 30 mins at 4 ºC/RT. This step was repeated two times, with 10 ml of solvent mix (1:1) and 15 ml of 1:2 ratio for 24 hours, and centrifuged to collect and pool the supernatant. The pooled supernatant was filtered using a nylon filter (0.2 μm) and dried at 37 º in a SpeedVac. The chloroform (100 μl) resuspended dried material was separated on a TLC plate coated with fluorescent indicator F254 (250 μm, silica, Merck) and mobile phase (petroleum ether: ethyl acetate; 98:2, v/v). The plate was visualised under short UV in the UV transilluminator and stained in an iodine chamber.

Mtb H37Rv cultures from the log phase were harvested for whole-cell lysate preparation. Briefly, the culture pellet, after centrifuging at 3,500 rpm for 10 minutes at room temperature, was heat-inactivated at 95 ºC for 15 minutes in a water bath, followed by incubation for 30 minutes at room temperature. The heat-inactivated bacteria were resuspended in lysis buffer (PBS containing 8 mM EDTA, 1X proteinase inhibitor, 0.05 mg/ml RNase, 0.2 μg/ml DNase), zirconia beads (100 mg, 0.1 mm) were added for bead beating using a Biospec Mini-Beadbeater-16. A set of 3 cycles at 3650 oscillations/minute with 30 s on/off cycles for six times with 30 s breaks on ice between each cycle was adopted. This preparation was centrifuged at 600 rcf for 5 minutes at 4ºC to remove unbroken cells, and the supernatant was further clarified by centrifuging at 10,000 rcf for 10 minutes at 4 ºC. To make a cell-free preparation, the lysate was passed through a nylon filter (0.2 μm), and the total protein was quantified using BCA assay.

### Aerosol infection of mice

The eu- and hyper-glycemic C57BL/6 mice were infected with low aerosol dose (100-120 CFUs) of Mtb H37Rv using a Madison chamber in the Tuberculosis Aerosol Challenge Facility (BSL-III) at the host institute (ICGEB, New Delhi). For this, Mtb H37Rv culture (mid-log phase, OD ∼ 0.6) was harvested by centrifuging at 3,500 rpm at room temperature for 10 minutes and subjected to single cell preparation by resuspending in PBS (1 ml) and passing through a 23-gauge needle followed by 26-gauge needle (5 times each). Mice were aerosol infected in the Madison chamber (Gilmont parameter: 35, Pressure gauze: 25, Magnehelic: 1, Photohelic: 7) so that a dose of 100-120 Mtb reaches to the mice lungs. On day 1, a subset of mice was sacrificed to monitor the mycobacterial load. At 21 days post-infection, mice of eu- and hyper-glycemic mice with their group-specific controls were sacrificed to monitor their lung mycobacterial load. The femur and tibia of these mice groups were used for harvesting bone marrow-derived macrophages. The lung homogenates were plated on 7H11 agar supplemented with OADC to enumerate the CFU by 4 weeks post-incubation in the humidified chamber.

### Cell isolation, culture and infection

Bone marrow (BM) cells were harvested from the femur and tibia. After RBC lysis, viability was monitored by trypan blue staining, and harvested BM cells were cultured in BMDM media containing DMEM, FBS (10%), and M-CSF (40 ng/ml, BioLegend). Using PBS-EDTA (5 mM), on day 7, BMDMs were harvested and seeded on a 6-well plate (4 million cells per well for RNA extraction and 2 million cells for cytokine estimation). After allowing the BMDMs to attach overnight, they were washed twice with PBS and incubated with fresh BMDM media. A set of harvested cells (2.0 million/mouse) were incubated with an antibody cocktail (live and dead stain near-IR, CD45-PerCP Cy5.5, CD11c-FITC, CD11b-APC, F4/80-PE) for 30 mins at RT and after fixing with paraformaldehyde (4%), flow cytometry data were acquired on LSR Fortessa X-20 (BD Biosciences) and analysed using FCS-express (Denovo software, version 6).

### H37Rv lysate stimulation on BMDMs

A subset of BMDMs harvested from all the study groups, except TB, was stimulated by incubating with Mtb H37Rv lysate (10 μg/ml protein equivalent) at 37 ºC in 5% CO_2_. Cells were harvested 24 hours post-stimulation using TRIzol reagent (Invitrogen) for RNA. Briefly, 0.2 ml chloroform was added to the lysed cells per 1 ml TRIzol and mixed by shaking and incubating for 2-3 minutes. The cells were centrifuged for 15 minutes at 12,000 rcf for 15 minutes at 4 ºC. The aqueous phase containing RNA was transferred to a new tube, and isopropanol (0.5 ml) was added. After 10 minutes of incubation at 4 ºC, the solution was centrifuged at 12,000 rcf for 10 minutes at 4 ºC. The pellet was resuspended in ethanol (1 ml, 75 %) and vortexed briefly, centrifuged at 7,500 rcf for 5 minutes at 4 ºC. The RNA pellet was air-dried and resuspended in nuclease-free water. RNA was quantified by Qubit fluorometer using a Broad range RNA quantification kit (Invitrogen), and RNA integrity (RIN) value was obtained from the TapeStation (Agilent Technologies) using a Broad range RNA kit. The library was prepared from RNA (0.5 μg/sample) and sequencing was carried out with Illumina Paired End chemistry and raw data files in fastq format were subsequently processed as explained in the following section. Due to the low RNA quality, samples extracted from the TB group of 24 hours post stimulation were excluded from the data acquisition.

### RNA-seq data processing and DEG analysis

RNA-seq data in fastq.gz format was subjected to adapter trimming using fastp v0.20.1 (options used -l 75) (link:www). Adapter trimmed reads were aligned on the prebuilt hisat2 genome index of Mus musculus mm10 (https://genome-idx.s3.amazonaws.com/hisat/mm10_genome.tar.gz) using hisat2 v2.2.1. The reads were sorted using samtools with default options. Alignment files (BAM) were assessed with Qualimap v2.3 (https://bitbucket.org/kokonech/qualimap/downloads/qualimap_v2.3.zip). De novo transcript assembly and quantitation were performed with StringTie v2.2 (https://github.com/gpertea/stringtie). The transcript count matrix was prepared using the prepDE.py script from the StringTie package. For PCA analysis, the counts were transformed using edgeR (options: Min. CPM ≥ 0.05 in at least RNAseq datasets; pseudo count = 4; missing values were treated as zero). For the determination of differentially expressed genes (DEGs), DeSeq2 was used. The genes with log_2_FC ≥ 2.0 with adjusted P-value < 0.05 were considered deregulated and subsequently used for pathway enrichment analysis. The pathway analysis was done using prefranked GSEA. PCA analysis, DEG determination, and pathway enrichment were performed on the iDEP 2.0 shiny-based platform (https://github.com/gexijin/idepGolem) (Supplementary Figure 2A).

Statistical Analysis: All data are presented as mean ± standard deviation (SD) values. Statistical analyses were performed using DeSeq2, edgeR, iDEP 2.0, GraphPad Prism (Version 8.4.2) and Microsoft Excel. Differences were considered statistically significant with log_2_FC ≥ 2.0 with adjusted P-value < 0.05(for transcriptomics data) and p<0.05 (for others) at 95% confidence.

### Online supplemental material

This manuscript contains supplementary information.

## Abbreviations

TB: (Tuberculosis)
DM: (Diabetes mellitus)
BM: (Bone marrow)
BMDM: (Bone marrow-derived macrophages)
OD: (Optical density)
CFU: (Colony forming units)
DMEM: (Dulbecco’s modified eagle medium)
GM-CSF: (Granulocyte monocyte-colony stimulating factor)
BCA assay: (Bicinchoninic assay)
TEAB: Triethylammonium bicarbonate buffer)
EDTA: (Ethylenediaminetetraacetic acid).

## Acknowledgements

Nidhi Yadav and Ashish Gupta received Junior and Senior Research Fellowships from the Department of Biotechnology (DBT), Government of India. Shweta Chaudhary received a Shyama Prasad Mukherji fellowship from the Council of Scientific and Industrial Research. Nikhil Bhalla is supported by the DBT-funded project (Grant ID: National Network Project of National Institute of Immunology, New Delhi -[40267]). We acknowledge the DBT, Government of India, for supporting activities through research grants and supporting the Tuberculosis Aerosol Challenge Facility (TACF) at ICGEB and ICGEB New Delhi for providing access to core support to RKN.

## Authorship Contributions

NY and RKN conceptualized and designed experiments. SC developed the diabetes induction model and performed the Mtb aerosol challenge experiment. NY and AG performed the BM isolation and in-vitro experiments; NB and NY analysed the data and prepared figures. Funds for this work were generated by RKN; NY, NB and RKN wrote the manuscript and revised it, incorporating the comments of all co-authors.

