## Supplemental Information for "Macrophages of diabetes and tuberculosis co-morbid conditions show perturbed differentiation and metabolic signatures"

**Supplementary figure 1**

**Supplementary Figure 1: Flow cytometry of BMDMs A. Representative gating strategy followed to monitor CD11c, CD11b and F4/80 expression on BMDMs B. Percentage of CD11c^+^, CD11b ^+^, F4/80 ^+^ cells in different groups. Abbreviations: BMDMs: Bone marrow-derived macrophages.**

**A**

**B**

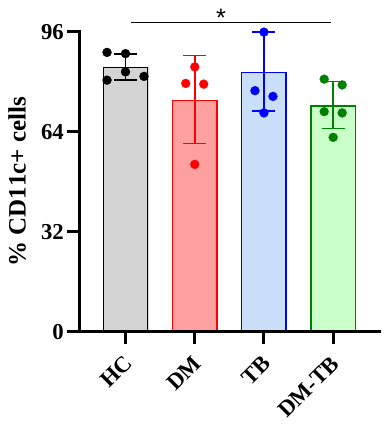

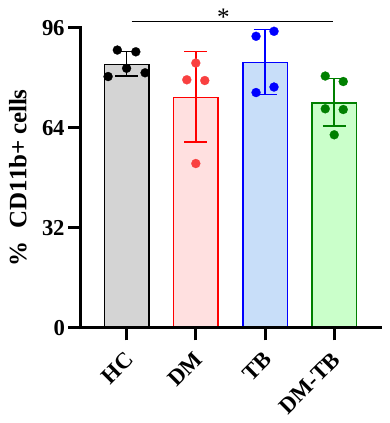

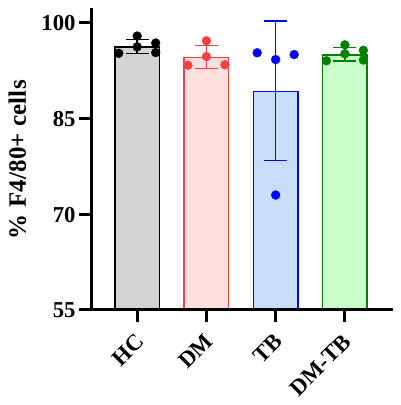

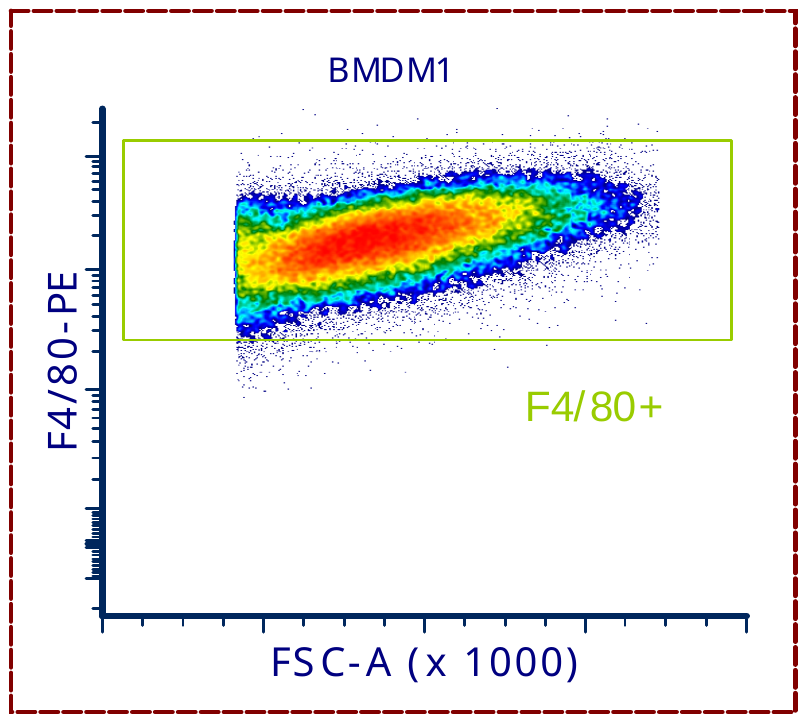

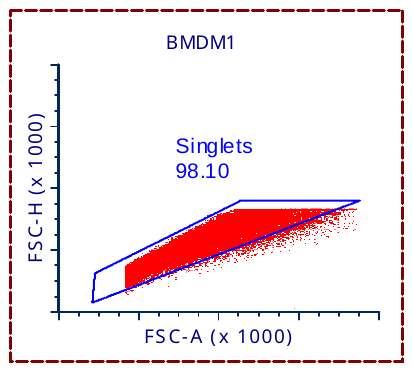

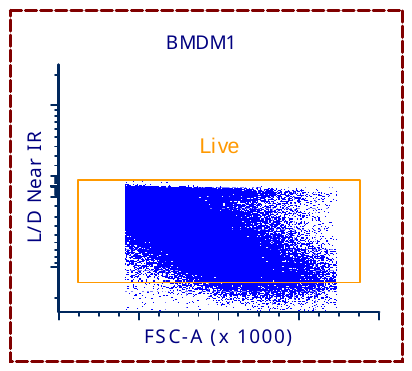

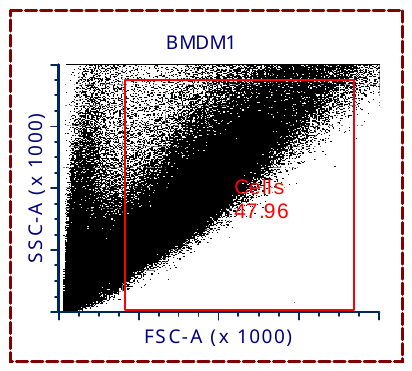

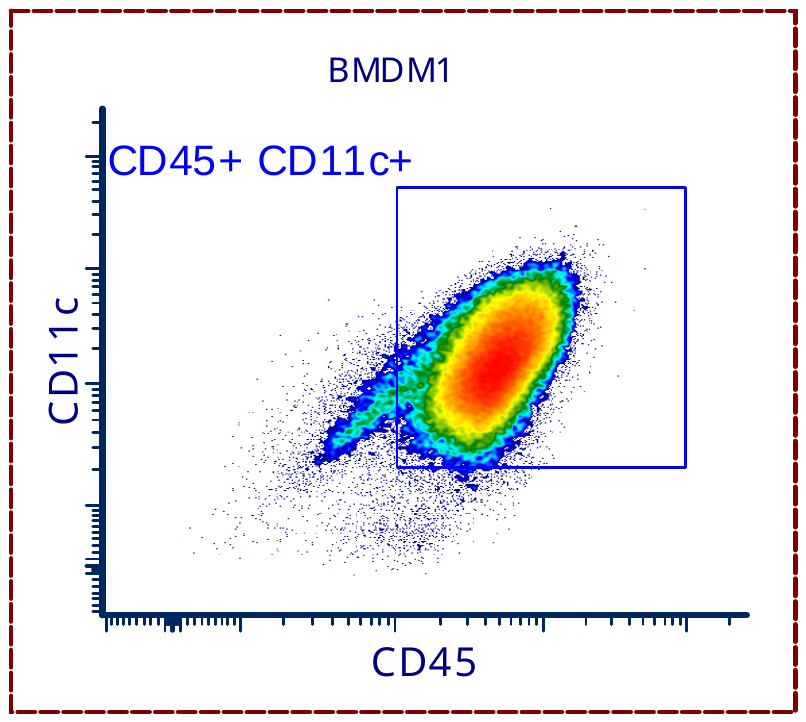

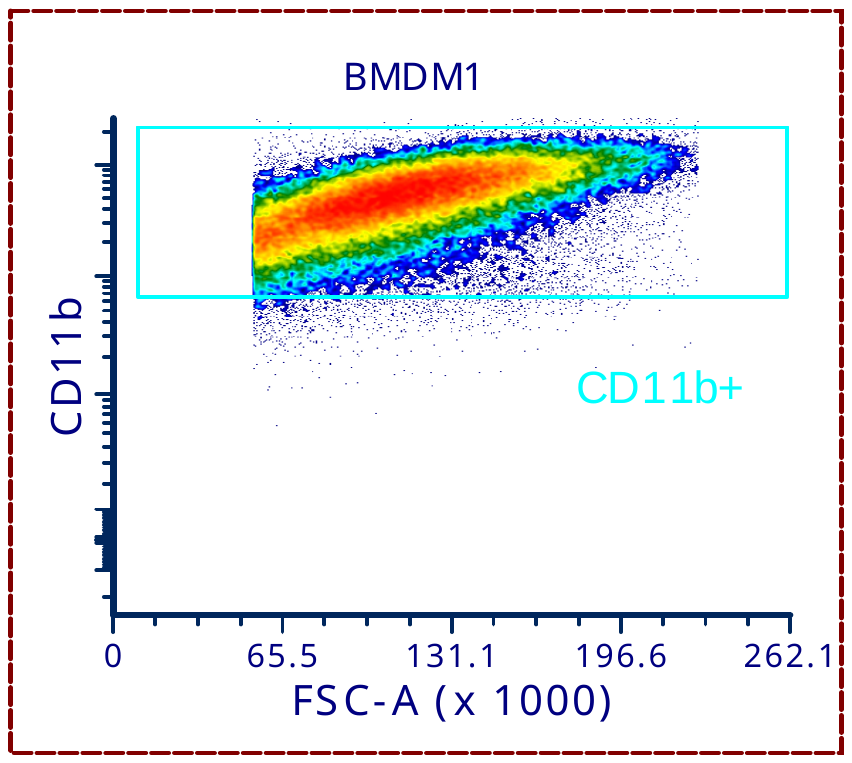

**Supplementary Figure 2: Other permutations showed interesting transcriptome differences. A, B and C: Volcano Plots showing DEGs between various groups highlighted with red and blue dots.**

Abbreviations: BMDMs: Bone marrow-derived macrophages, DEGs: Deregulated genes, DM: Diabetic, HC: Healthy controls, TB: Tuberculosis, DM-TB Diabetes-tuberculosis comorbidity.

**Supplementary figure 2**

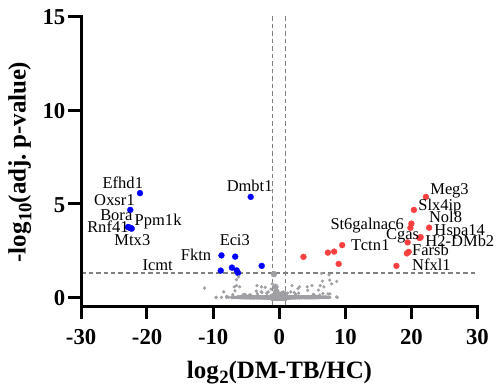

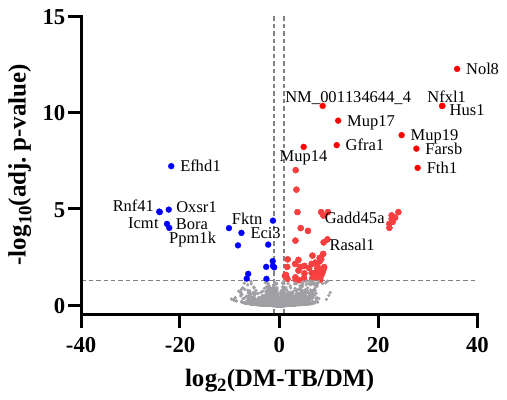

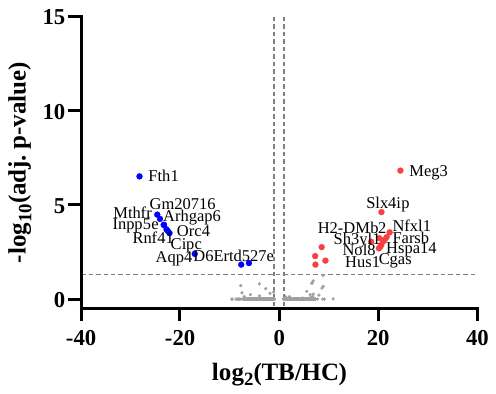

**C**

**A**

**B**
